# Biflavonoids can potentially inhibit amyloid beta internalization to mitigate its cytotoxic events

**DOI:** 10.1101/2025.01.16.633335

**Authors:** Md. Aminul Haque, Md. Selim Hossain, Il Seon Park

**Author notes:** Corresponding authors: (MAH); (ISP).

## Abstract

Amyloid-β-42 (Aβ) peptides are key contributors to Alzheimer’s disease (AD), with recent studies highlighting a strong link between Aβ toxicity and its internalization into cells. Targeting the internalization of Aβ using naturally occurring small molecules could represent a promising strategy for developing therapeutic interventions. In this study, we investigated the effects of biflavonoids on inhibiting the internalization of Aβ and its impact on cellular processes that lead to cell death. Various biochemical techniques were employed to assess the ability of biflavonoids to prevent Aβ42 uptake and to elucidate the underlying mechanisms of their inhibitory action. Our findings revealed that biflavonoids exert a dose-dependent inhibitory effect on Aβ toxicity. The cytoprotective effects of biflavonoids were primarily attributed to their ability to block Aβ internalization, as confirmed through confocal microscopy and validated by western blot analysis. By preventing Aβ entry into cells, biflavonoids also inhibited Aβ-induced lamin fragmentation and caspase activation, both of which are critical steps in Aβ-mediated cytotoxicity. Furthermore, we explored the influence of biflavonoids on the aggregation process of Aβ, including fibril, oligomer, and β-sheet formation, and found that biflavonoids effectively inhibited these conformational changes. Based on these results, we propose that biflavonoids suppress Aβ cytotoxicity by blocking its conformational changes and subsequent internalization, thereby highlighting their potential as anti-amyloidogenic therapeutic agents.

## Introduction

Alzheimer’s disease, also referred to as senile dementia, is the most prevalent form of dementia, characterized as a neurodegenerative disorder that predominantly affects the elderly. The incidence of AD significantly increases with age, with studies indicating that up to 50% of individuals aged 85 and older are affected by this condition [1]. The number of people with AD diagnoses is predicted to increase in line with the aging of the world’s population. Amyloid-β peptides, which are short chains of 39–43 amino acids and are central to the pathophysiology of AD, are involved in the disease’s pathogenesis [2]. These peptides are synthesized by the proteolytic cleavage of amyloid precursor proteins (APP) by the enzymes α-, β-, and γ-secretases [3]. Amyloid-β in its monomeric form exhibits a tendency to misfold and take on structures rich in β-sheets. These transitional structures have a tendency to aggregate, resulting in the formation of oligomers, protofibrils, and ultimately mature fibrils [4]. These aggregates accumulate in the brain’s parenchyma and cerebral blood vessels, contributing to plaque formation, which is an essential component of AD pathology [5]. Remarkably, studies that previously concentrated on deposited fibrils have now turned their attention to soluble oligomers and protofibrils, which are thought to be more neurotoxic. It is believed that these soluble forms of amyloid-β are essential for the development and progression of AD, resulting in synaptic dysfunction and neuronal damage prior to the external appearance of fibrillar plaques [6–11].

In neurons, the build-up of amyloid-β-42 has been suggested as an important factor for the pathogenesis of Alzheimer’s disease. Moreover, it is notable that prolonged expression of human amyloid precursor protein (APP) in rat cortical neurons resulted in apoptosis, which is a type of programmed cell death [12]. The investigation further distinguished between the effects of different amyloid-β peptides and found that whereas the extracellular amyloid-β-40 produced by processing APP did not cause neuronal death, the intracellular accumulation of amyloid-β-42 resulting from full-length APP expression led to apoptosis. This implies that when amyloid-β-42 is present inside cells, it has a special tendency to cause neuronal damage. Furthermore, it has been noted that neurons can internalize the oligomeric forms of amyloid-β via endocytosis [13], however the less toxic fibrillar forms are typically not taken up by cells in this way [14]. The greater cytotoxicity of oligomers in comparison to other forms of amyloid-β could potentially be explained by their ability to enter neurons. These oligomers’ intracellular location is especially troublesome since it could intensify their harmful effects. Amyloid-β peptides can be potentially dangerous both inside and outside of cells, however the degree of cytotoxicity greatly depends on its location [15]. Our previous study demonstrated that the cytotoxicity is more reliant on the intracellular presence of the peptide [15]; therefore, preventing its entry into the cell could be an effective strategy to reduce Aβ-induced toxicity.

The most prevalent kind of polyphenols in the human diet are flavonoids, a broad and diverse type of polyphenolic chemicals that are abundant in plants [16]. These compounds are well-known for their multiple health benefits, which include antibacterial, anti-allergic, anti-cancer, and antioxidant properties [17–19]. Notably, flavonoids have also been associated with a reduced risk of age-related dementia [20]. A higher dietary intake of flavonoids, especially flavonols, is linked to a reduced prevalence of dementia, according to epidemiological research carried out in 23 developed countries [21]. Supporting this, a large cohort study involving a random sample of 1,367 individuals aged 65 and older found an inverse relationship between the consumption of flavonoid-rich foods and the risk of developing dementia [20]. Further research has focused on specific flavonoid-rich extracts, such as those derived from Ginkgo biloba, namely HE208 [22] and and EGb761 [23]. These studies have demonstrated that flavonoid molecules are instrumental in exerting anti-amyloidogenic and anti-apoptotic effects on neural cells. Furthermore, it has been found that a number of isolated flavonoids can efficiently prevent the development of amyloid-β oligomers as well as fibrils, and reduce the cytotoxic effects of amyloid-β peptides [24–28]. However, the connection between the protective properties of flavonoids and their structure is complicated. While some polyphenols can inhibit the fibrillogenesis of amyloid-β peptides, they may not necessarily prevent amyloid-β-induced cytotoxicity. Conversely, some flavonoids provide cytoprotection against amyloid-β without possessing antifibrillogenic properties [29, 30]. This implies that the mechanisms behind flavonoids’ anti-cytotoxic and anti-amyloidogenic properties are not completely known and may differ based on the particular flavonoid molecule and its interactions with amyloid-β peptides. Considering the complex nature of these interactions, more investigation is necessary to determine the specific role of flavonoids that can play in the prevention or treatment of neurodegenerative conditions like Alzheimer’s disease. In the current study, we explored the association of bioflavonoids impact on the prevention of peptide’s entry into the cell with its cytotoxicity.

## Materials and Methods

### Materials

Taiwaniaflavone (TF) and amentoflavone (AF) were isolated following the methodologies detailed in previous research [32]. Fetal bovine serum (FBS) was procured from Atlas Biologicals Inc. (Fort Collins, Colorado, USA). Dulbecco’s Modified Eagle Medium (DMEM) and the antibiotic solution penicillin/streptomycin (P/S) were obtained from Welgene (Gyeongsangbuk-do, Korea). Urea and phosphate-buffered saline (PBS) were purchased from Georgia Chemical & Equipment Company Inc. (Norcross, Georgia, USA) and Ameresco (Framingham, Massachusetts, USA), respectively. Methanol and ethanol were supplied by OCI Company Ltd. (Seoul, Korea). Additionally, isopropyl β-D-1-thiogalactopyranoside (IPTG) was sourced from Bioneer (Daejon, Korea). All other reagents were collected from Sigma-Aldrich (St. Louis, Missouri, USA), unless otherwise stated.

### Preparation of Aβ peptides

Aβ peptides in form of fusion proteins were initially expressed in E. coli, following established protocols [33]. After expression, the peptides were purified and then 100% 1,1,1,3,3,3-hexafluoro-2-propanol was used to dissolve it. The dissolved peptides were then dried under a nitrogen flow, followed by an additional drying phase under vacuum for 30 minutes. The resulting peptide aliquots were preserved at –20°C. The peptides (2 mg/mL) were dissolved in 0.1% NH4OH before using. For better dissolution, it was then sonicated at 4°C for 10 minutes. The sonicated peptides were subsequently diluted to the desired working concentration using phosphate-buffered saline (PBS) or cell culture media, depending on the experimental requirements. For the formation of oligomers and fibrils, freshly prepared Aβ peptides were used. To promote oligomer formation, the peptide solution was mixed thoroughly into serum-free DMEM (without phenol red) and incubated at 4°C for 24 hours at a concentration of 100 µM. After that, the solution was centrifuged at 16,000 × g for 15 minutes to separate the supernatant, which was then diluted to a final concentration of 20 µM using serum-free DMEM. For fibril preparation, a 20 µM peptide solution in PBS was prepared and incubated at 37°C for 24 hours.

### Cell Culture and Cell Viability Assay

Human epithelial HeLa cells were cultured following previously explained procedure [34]. In 96-well plates (Nunc, Roskilde, Denmark), the cells (15,000/well) were seeded and cultured for 24 hours for the cell viability assessment. After a further 12 hours of serum deprivation, the cells were treated in accordance with the treatment plan. The MTT reduction test was used to determine the viability of the cells [35], where 20 µL solution (5 mg/mL MTT in PBS) was added to each well. 100 µL of solubilization buffer [20% sodium dodecyl sulfate (SDS) solution in 50% (v/v) N, N-dimethylformamide (DMF) (pH 4.7)] was added after 2 hours of incubation and the mixture was incubated for additional 12–16 hours. At 570 nm, absorbance was measured with a microplate reader (KisanBio, Seoul, Korea). AlamarBlue assay was also utilized to assess the cytotoxicity [36], where in each well 10 µL of alamarBlue solution (Life Technologies, Inc., Carlsbad, California, USA) was added directly and kept for 4-16 hours. Gemini-XS microplate spectrofluorometer (Molecular Devices, San Jose, California, USA) was used to measure the fluorescence at excitation and emission wavelengths of 560 and 590 nm, respectively.

### Observing the cellular localization of Aβ peptide via confocal microscopy

In a 12-well plate, cells (1 × 105) were plated and incubated for 24 hours, followed by 12 hours serum deprivation at 37°C where serum-free medium was used. Cells were treated and fixed in methanol at –20°C and permeabilized with 0.3% Triton X-100. 0.1% BSA (Bovine Serum Albumin) was used for blocking. Following overnight blocking, mouse monoclonal anti-Aβ antibody 6E10 (BioLegend, San Diego, California, USA) or rabbit polyclonal anti-caspase-9 (p10) antibody (Santa Cruz Biotechnology, Santa Cruz, California, USA) was added to each sample and kept overnight at 4°C. Alexa-Fluor-546-TRITC-conjugated goat anti-rabbit IgG and Alexa-Fluor-488-FITC-conjugated goat anti-mouse IgG antibodies (dilution, 1:200, Invitrogen, Waltham, Massachusetts, USA) were added after washing with PBS, kept for 2 hours at room temperature, and then again washing was done with PBS. DAPI in Vectashield mounting medium (Vector Laboratories, Burlingame, California, USA) was used to stain the nuclei. Carl Zeiss LSM510 microscope (Oberkochen, Germany) with the manufacturer’s software (LSM 510) was employed to take the confocal images, as previously described [37]. Four individual variable pinholes (97 µM) of 1.0 airy units per confocal channel were used with a resolution of 2048 × 2048 pixels. Plan apochromat 63 × 1.4 oil immersion objective were applied to focus the cells and cells were projected in a single plane.

### Detection of Aβ internalization and lamin fragmentation via immunoblotting

In a 60 × 15 mm cell culture dish, cells (4 × 105) were seeded and cultured at 37°C for 24 hours to ensure proper cell attachment and growth. Following this incubation, the cells were subjected to serum deprivation for an additional 12 hours. After treatment, the cells were harvested and washed twice with ice-cold PBS. To ensure the removal of surface-bound Aβ peptide, 0.1% Triton X-100 was added to the PBS during the washing steps. This ensures that any measured internalization of Aβ peptide is accurate and not due to residual surface-bound peptide. After harvesting the cells, they were lysed in a lysis buffer [50 mM Tris-HCl (pH 8.0), 150 mM NaCl, 1% Triton X-100, 5 mM EDTA, 5 mM EGTA, 1 mM PMSF, 10 µg/mL leupeptin, 2 µg/mL pepstatin A, and 2 µg/mL aprotinin]. To facilitate the complete cell lysis, cells were kept on ice for an additional 20 minutes. Following lysis, the cell lysates were centrifuged at 4°C at a speed of 18,000 × g for 15 minutes. This high-speed centrifugation step is necessary to pellet cell debris and isolate the supernatant containing soluble proteins. Upon careful collection of the supernatant, protein concentration was measured by the Bradford assay to ensure equal loading of samples for subsequent analysis. Equal amounts of protein from each sample were subjected to 12-15% SDS-polyacrylamide gel electrophoresis (SDS-PAGE), following the protocol previously described [38]. Proteins were transferred to a polyvinylidene fluoride (PVDF) membrane following electrophoresis. The membrane was immunoprobed with mouse monoclonal anti-Aβ 6E10 (BioLegend, San Diego, California, USA), lamin B, and β-actin antibodies (Santa Cruz Biotechnology, Santa Cruz, California, USA) followed by horseradish peroxidase (HRP) conjugated anti-mouse secondary antibodies (Santa Cruz Biotechnology, Santa Cruz, California, USA) [37]. The immunoblots were visualized using WESTER ɳC ULTRA (Cyanagen, Bologna, Italy), a highly sensitive chemiluminescent substrate. This allows for the detection and quantification of the target proteins, providing insights into the expression and localization of the Aβ peptide and other proteins of interest under the experimental conditions.

### Assessment of caspase activity

In 96-well plate, cells (2 × 104) were plated and incubated at 37°C for 24 hours to allow for adherence and initial growth. Following this incubation period, the cells were serum-starved for an additional 12 hours and treated according to treatment plan. After treatment, two times washing were done using ice-cold PBS to remove any residual media and treatment compounds, ensuring that subsequent assays were not influenced by external factors. Following the washing steps, cells were lysed using 40 μL of cell lysis buffer [20 mM HEPES-NaOH (pH 7.0), 1 mM EDTA, 1 mM EGTA, 20 mM NaCl, 0.25% Triton X-100, 1 mM DTT, 1 mM PMSF, 10 µg/mL leupeptin, 5 µg/mL pepstatin A, 2 µg/mL aprotinin, and 25 µg/mL N-acetyl-Leu-Leu-Norleucinal]. The buffer components were carefully selected to ensure efficient cell lysis and preservation of protein integrity, while protease inhibitors were included to prevent protein degradation. The cell lysates were kept on ice for 20 minutes to facilitate complete lysis and protein extraction. Following lysis, 50 μL of caspase assay buffer [20 mM HEPES-NaOH (pH 7.0), 20 mM NaCl, 1.5 mM MgCl2, 1 mM EDTA, 1 mM EGTA, and 10 mM DTT] was added to each well. Caspase-3 (DEVDase) activity was initiated by adding 10 μM Ac-DEVD-AMC (AG Scientific, Inc., San Diego, California, USA) to each well. The reaction was conducted at 30°C. The release of AMC (7-amino-4-methylcoumarin), a fluorescent product, was monitored for 1 hour at 5-minute intervals. The fluorescence was measured using a Gemini-XS microplate spectrofluorometer (Molecular Devices, San Jose, California, USA) with excitation and emission wavelengths set at 360 nm and 480 nm, respectively. This approach allows for the continuous monitoring of caspase-3 activity in real time, providing a kinetic profile of enzyme activity under the experimental conditions. The fluorescence data were collected and analyzed to determine the rate of AMC release, which directly correlates with caspase-3 activity. This detailed methodology ensures accurate and reproducible measurements of caspase-3 activity, facilitating a better understanding of the cellular response to the treatments applied in this study, as previously described in our protocols [39].

### Transmission electron microscopy (TEM)

To prepare the samples for transmission electron microscopy (TEM) analysis, a 5 µL aliquot was carefully deposited onto a Formvar-coated 200-mesh nickel grid, which was sourced from SPI Supplies (West Chester, Pennsylvania, USA) and kept for 5 minutes and then excess solution was removed using filter paper. The grid was then washed three times with distilled. Following the washing, the grid was subjected to negative staining with a 2% uranyl acetate solution for 1 minute. After staining, the grids were thoroughly examined under a transmission electron microscope (H-7600, Hitachi, Tokyo, Japan). The TEM was operated at an accelerating voltage of 80 kV and observations were conducted at a magnification of 40,000× [40, 41].

### Aβ Fibrillogenesis Assessment by Th-T Assay

To assess the fibrillogenesis of the peptide solution, 300 µL of a 20 µM peptide solution prepared in phosphate-buffered saline (PBS) was incubated at 37°C. From this solution, a 20 µL aliquot was carefully taken and mixed with 80 µL of a freshly prepared 5 µM thioflavin T (ThT) (Bioneer, Daejon, Korea) solution in PBS. The mixture of peptide and ThT was then subjected to fluorescence measurement which was carried out using a Gemini-XS microplate spectrofluorometer (Molecular Devices, San Jose, California, USA) [6]. The spectrofluorometer was set to detect fluorescence at an excitation wavelength of 445 nm and an emission wavelength of 490 nm. This method, as previously established in the literature [10], allows for the quantification of fibril formation by monitoring the increase in fluorescence intensity over time.

### Circular Dichroism Spectroscopy

To investigate the secondary structure of the Aβ peptide in solution, circular dichroism (CD) spectra were recorded using a Jasco spectropolarimeter (Tokyo, Japan) as previously described [42]. The measurements were conducted in a 1 mm pathlength cuvette. The spectra were collected at 0.5 nm intervals with a resolution of 1 nm across the wavelength range of 190 to 250 nm at 25°C. The CD spectra were recorded at a scan rate of 50 nm/min. For each sample, five individual scans were performed and then averaged. For experiments involving the incubation of Aβ with biflavonoids, spectra were obtained by subtracting the background spectra of the buffer.

## Results

### Cytoprotection against Aβ induced toxicity

In this study, we investigated the cytotoxic effects of various Aβ peptide preparations, specifically monomeric Aβ (mAβ), oligomeric Aβ (oAβ), and fibrillar Aβ (fAβ), using HeLa cells. To assess cell viability, we utilized both MTT and alamarBlue assays, which provided comparable results, ensuring the reliability of our observations. Among the three types of Aβ preparations, fAβ exhibited the lowest level of cytotoxicity, as depicted in Figures 1A and 1B. This finding led us to exclude fAβ from further detailed exploration, as it did not significantly impact cell viability compared to the other forms.

**Fig 1.**
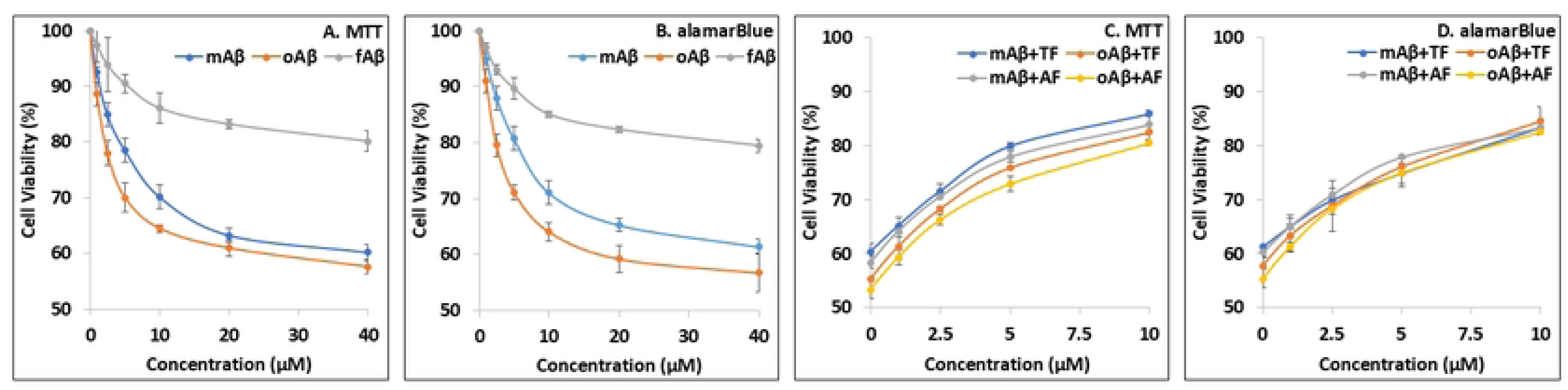
Cytotoxicity of Aβ in presence and absence of biflavonoids where HeLa cells were used. (A, B) Cell viability assessment of different types Aβ at different indicated concentrations by MTT assay and alamarBlue assay. (C, D) Dose dependent cytoprotection of biflavonoids against different types of Aβ (20 µM) using MTT assay and alamarBlue assay. Results are expressed as the mean ± standard deviation of values from three independent experiments.

Quantitatively, we observed that 20 µM concentrations of mAβ and oAβ resulted in approximately 35% and 40% cell death, respectively. These results highlight the differential toxicity of Aβ forms, with oAβ being the most cytotoxic. This observation is consistent with previous studies that have demonstrated the heightened toxicity of oligomeric forms of Aβ due to their higher propensity to disrupt cellular processes.

Following the initial assessment of Aβ-induced cytotoxicity, we shifted our focus to evaluating the cytoprotective effects of specific biflavonoids, namely TF and AF, as illustrated in Figures 1C and 1D. Both TF and AF exhibited a dose-dependent inhibition of cell death across both mAβ and oAβ treatments. Notably, at a concentration of 10 µM, these biflavonoids significantly enhanced cell viability, increasing it from a baseline of 55-60% to an improved range of 80-85%. This marked increase in cell survival indicates the potent protective effects of TF and AF against Aβ-induced cytotoxicity.

To further understand the mechanism behind this protection, we explored whether biflavonoids could prevent the internalization of Aβ peptides into cells. This line of investigation was crucial, as the entry of Aβ into cells is a key factor in initiating cytotoxic pathways. Our results suggest that the ability of TF and AF to inhibit Aβ internalization and hence halt the activation of harmful downstream pathways may contribute to their protective effects which underscores the potential of biflavonoids as therapeutic agents in reducing Aβ-induced cellular damage.

### Inhibition of Aβ internalization into cells thus inhibiting cytotoxic events

The impact of biflavonoids on the internalization of Aβ peptides was investigated using confocal microscopy (Figure 2A) and western blotting (Figure 2B). We specifically focused on the oligomeric form of Aβ (oAβ) because of its superior ability to penetrate cells compared to other forms of Aβ. To trace the internalization process, we stained the cytoplasm using caspase-9, and it has been found that oAβ effectively entered the cells (Figure 2A).

**Fig 2.**
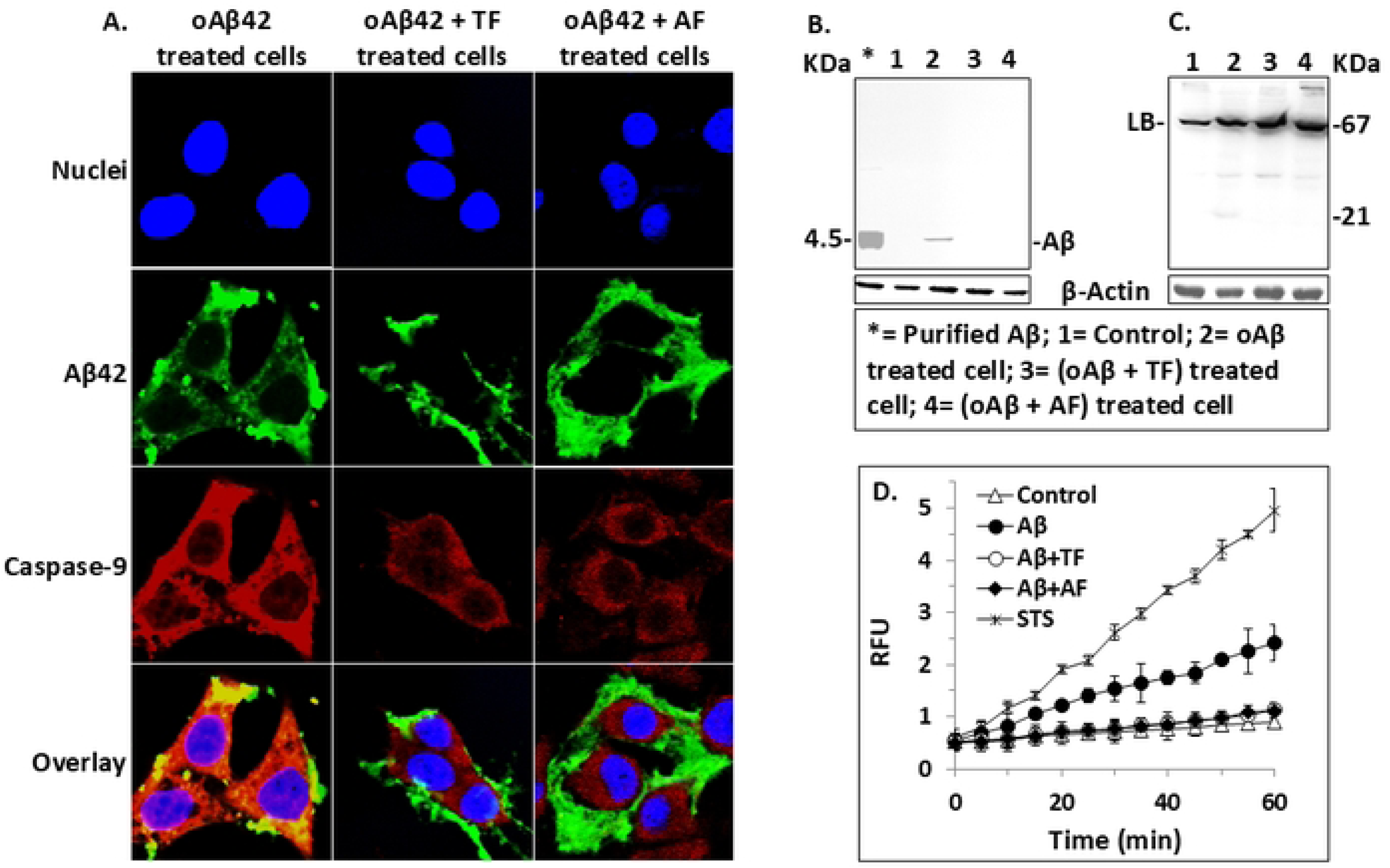
Effect of biflavonoids (10µM) on Aβ internalization. HeLa cells were treated with 20µM Aβ in presence and absence of biflavonoids for 24 hours. (A) Confocal microscope images of Aβ and caspase-9 were taken where monoclonal mouse anti-Aβ (6E10) and polyclonal rabbit anti-caspase-9 (p10) antibodies were applied to identify Aβ (green) and caspase-9 (red) which were visualized by using secondary goat anti-mouse IgG and goat anti-rabbit IgG antibodies, respectively. Nuclei were stained with DAPI (blue). Caspase-9 was used to locate the cytoplasm helped to monitor the location of Aβ also. Yellow spots indicate the interaction of caspase-9 and Aβ. In presence of biflavonoids, the internalization of Aβ was prevented. (B) Inhibition of Aβ internalization was measured using western blotting where mouse monoclonal anti-Aβ 6E10 and HRP conjugated anti-mouse secondary antibody were used. Cells without any treatment were used as negative control and the purified Aβ was used as positive control. (C) Prevention of caspase independent laminB fragmentation was monitored by western blotting where mouse monoclonal anti-laminB and HRP conjugated anti-mouse secondary antibody were used. Cells without any treatment were used as control. (D) DEVDase activity was measured with 10µM ac-DEVD-AMC substrate in HeLa cells treated with 20 µM Aβ with or without biflavonoids (10 µM). Staurosporine (STS)-treated (0.5µM for 6 h) cells were used as a positive control. Negative controls were the cells which were not treated with the peptide or STS. RFU indicates relative fluorescence unit. The results are the mean ± standard deviation of three independent experiments.

The impact of biflavonoids, specifically TF and AF, on the internalization of oAβ was then assessed. Figure 2A demonstrated the remarkable capacity of both substances to prevent the intracellular accumulation of oAβ. This suggests that TF and AF could potentially reduce the cytotoxic effects of oAβ by blocking its entry into the cells, thereby preventing it from initiating intracellular pathways leading to cell death. Later western blotting was employed to further validate the impact of these bioflavonoids on oAβ internalization. In oAβ treated cells, the peptide was readily detected upon cell lysis, confirming successful internalization. However, when cells were treated with oAβ in presence of TF and AF, the oAβ peptide was not detectable (Figure 2B, S1), indicating that these biflavonoids effectively prevented oAβ from entering the cells.

Combined results of western blotting and confocal microscopy indicated that TF and AF are potent inhibitors of Aβ internalization. This blockage is critical, as the internalization of Aβ is a prerequisite for its cytotoxicity [15]. In line with this, we examined subsequent intracellular events typically associated with Aβ toxicity, such as lamin fragmentation and caspase activation. Our data showed that biflavonoids not only prevented Aβ entry but also effectively inhibited lamin fragmentation (Figure 2C, S1) and caspase activation (Figure 2D). This combined inhibition further demonstrates the potential of TF and AF to mitigate critical intracellular processes that lead to cell death.

To investigate the effects of bioflavonoids on cells following Aβ internalization, cells were first incubated with various Aβ preparations for 24 hours, after which bioflavonoids were introduced. The initial assessment focused on cell viability, which revealed no significant inhibition of cytotoxicity (Figure 3A & 3B). As a result, effect on caspase activation and lamin fragmentation, were not further explored. These findings clearly indicate that once the Aβ peptides have entered the cells, bioflavonoids are unable to mitigate the ensuing cytotoxic effects.

**Fig 3.**
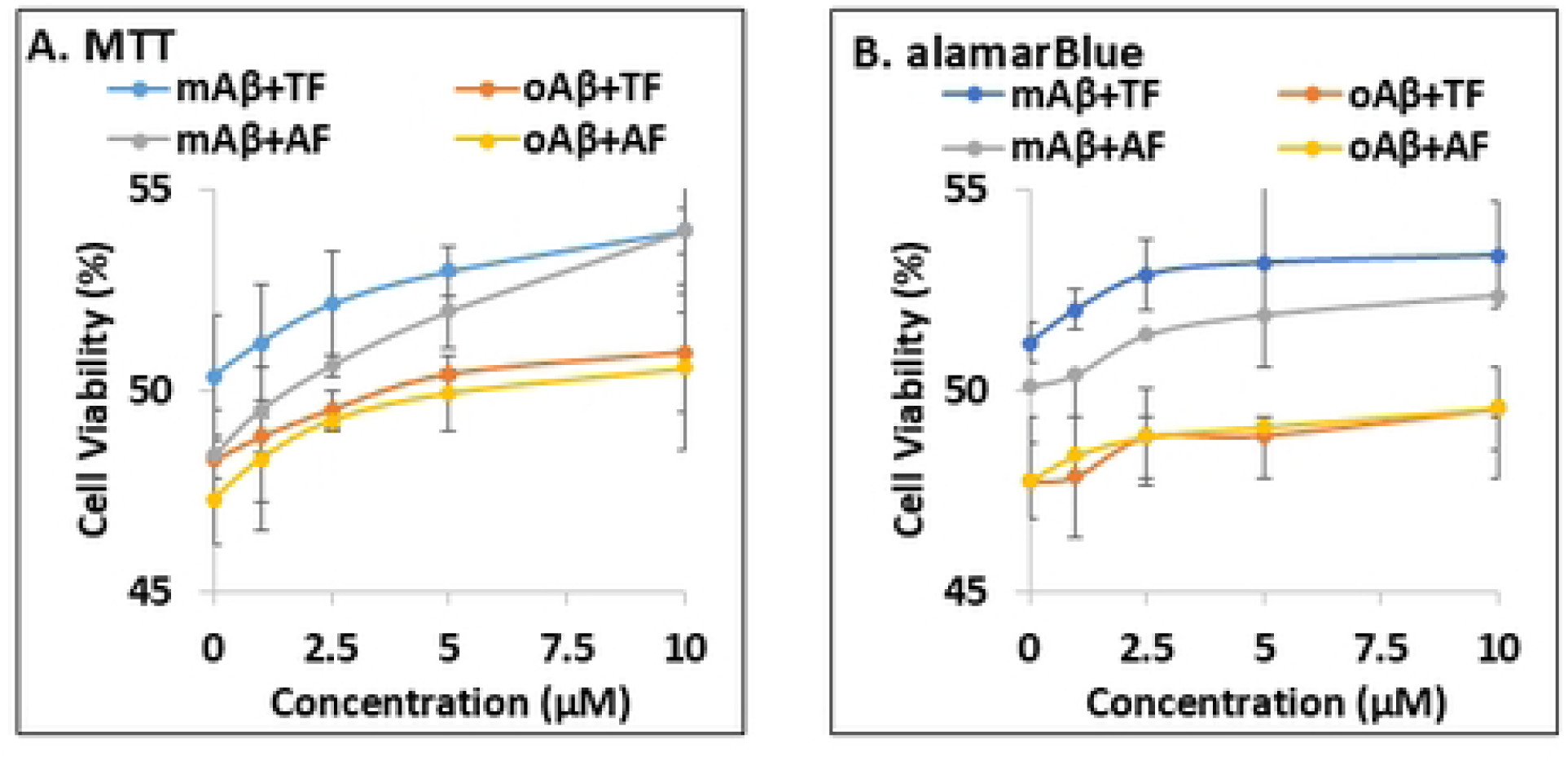
Cytoprotection ability of biflavonoids on cells following treatment with different types of Aβ. Initially cells were treated with 20µM mAβ and oAβ for 24 h and then bioflavonoids were introduced at different concentrations. Using MTT assay (A) and alamarBlue assay (B), cell viability was assessed. Results are expressed as the mean ± standard deviation of values from three independent experiments.

In summary, our findings highlight the significant role of biflavonoids in preventing the internalization of Aβ and the subsequent activation of harmful cellular pathways. By blocking Aβ entry and inhibiting key intracellular events such as lamin fragmentation and caspase activation, TF and AF emerge as promising agents for counteracting Aβ-induced cytotoxicity.

### Alteration of biophysical properties of Aβ peptides to prevent its cytotoxicity

Subsequently, our study delved into understanding the mechanisms by which biflavonoids inhibit the entry of Aβ peptides into cells. To clarify this mechanism, we investigated the effects of TF and AF on the key structural transformations of Aβ peptides like secondary β-sheets, oligomers, and fibrils formation. These structural forms are crucial in the progression of Aβ-induced toxicity.

Our results revealed that TF and AF significantly inhibited the formation of both oligomers and fibrils, as shown in Figure 4A. Oligomerization and fibrillogenesis are essential steps in the aggregation of Aβ peptides into toxic species. By inhibiting these processes, TF and AF reduce the potential of Aβ to aggregate into harmful forms. Moreover, secondary β-sheet formation which is considered as a critical structural transition necessary for the peptide to adopt its toxic conformation, was also markedly suppressed by both biflavonoids (Figure 4B).

**Fig 4.**
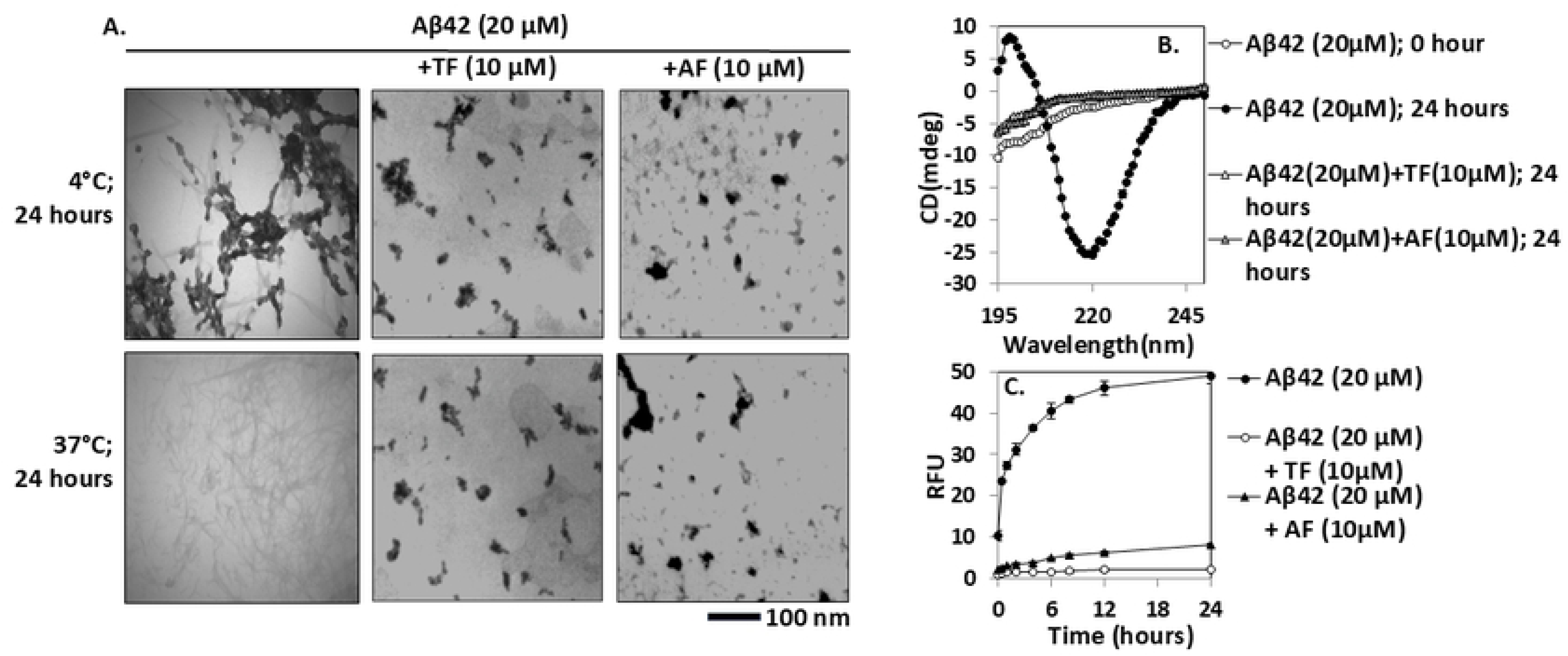
Effect of biflavonoids on biophysical properties of Aβ peptide. (A) Oligomer and fibril formation were assessed by collecting images with TEM at 40000×. Scale bars represent 100 nm in TEM images. (B) CD spectra analysis of 20 µM freshly prepared Aβ42 sample and incubated Aβ42 samples for 24 hours at 37°C with or without biflavonoids (10 µM). (C) Fibrillogenesis assessment by ThT-fluorescence assay using 20 µM Aβ42 peptides with or without biflavonoids (10 µM). RFU represents relative fluorescence unit. Triplicate experiments were performed and standard deviations were indicated as bars.

To further validate these observations, we employed the Thioflavin-T (Th-T) assay to assess fibrillogenesis, the process by which Aβ peptides form mature fibrils. The results from this assay confirmed that TF and AF significantly inhibited fibrillogenesis (Figure 4C), demonstrating their effectiveness in preventing Aβ aggregation.

Collectively, these findings suggest that biflavonoids, by disrupting key structural transitions like β-sheet formation, oligomerization, and fibrillogenesis, prevent the development of Aβ species capable of penetrating and damaging cells. This ability to interfere with the structural integrity of Aβ highlights the therapeutic potential of biflavonoids in mitigating the cytotoxic effects associated with Aβ aggregation, thereby offering promising avenues for developing treatments for diseases such as Alzheimer’s.

## Discussion

Flavonoids are well-documented for their neuroprotective properties in various conditions, including Alzheimer’s and Parkinson’s diseases. Their cytoprotective effects can be categorized into three key mechanisms: antioxidation, inhibition of Aβ aggregation, and prevention of Aβ-induced cell death [31]. However, the relationship between the anti-amyloidogenic properties and the anti-cytotoxic effects of these flavonoids is still not well understood. In this study, we concentrated on investigating how biflavonoids influence the internalization of Aβ peptides and, consequently, their cytotoxic effects.

Our findings revealed that the presence of various biflavonoids significantly reduced the cytotoxic effects of different forms of Aβ. To further understand the protective mechanisms of biflavonoids, their effects on Aβ internalization, confocal microscopy and western blotting were used where oligomeric form of Aβ was used due to its higher cell penetrability. Results showed that oAβ effectively penetrated cells, which is crucial for its cytotoxicity. However, both TF and AF significantly inhibited Aβ internalization, as evidenced by the absence of detectable Aβ in treated cells with biflavonoids (Figure 2A & 2B). These results emphasize the significance of biflavonoids in blocking Aβ entry into cells, which decreases Aβ’s cytotoxic effects. They also prevented caspase activation and caspase-independent lamin cleavage. These results collectively suggest that by blocking Aβ internalization, biflavonoids hinder the activation of caspases and prevent caspase independent lamin cleavage, which are both critical cellular events associated with Aβ-induced cytotoxicity. Consequently, the absence of Aβ within the cells disrupts the pathways leading to cell death.

Our subsequent investigations demonstrated that biflavonoids effectively inhibited the formation of secondary β-sheets, oligomers, and fibrils. The Thioflavin-T (Th-T) assay confirmed that biflavonoids significantly hindered fibrillogenesis, a key process in the aggregation of Aβ. By disrupting these structural transitions, biflavonoids prevent the formation of Aβ species capable of penetrating cells and causing toxicity. By preventing the aggregation of Aβ into these toxic forms, biflavonoids disrupt the pathways leading to cellular damage, thereby mitigating its harmful effects. This indicates that the structural modulation of Aβ by biflavonoids plays a crucial role in their protective function against Aβ-induced cytotoxicity by preventing its internalization. This approach allowed us to explore the potential therapeutic uses of biflavonoids in treating neurodegenerative disorders like Alzheimer’s disease as well as its wider therapeutic implications in preventing Aβ-induced cytotoxicity.

## Conclusion

In summary, the combined results from this study provide compelling evidence that biflavonoids offer substantial protection against Aβ-induced cytotoxicity. They achieve this by inhibiting Aβ internalization, preventing critical intracellular events, and disrupting the formation of toxic Aβ species. These findings suggest that biflavonoids could be promising therapeutic agents for neurodegenerative diseases characterized by Aβ aggregation, such as Alzheimer’s disease. Further research into their mechanisms and potential clinical applications is warranted to fully realize their therapeutic potential.

## Supporting information

**S1_raw_images. Full length blot image of Figure 2B & 2C**.

## Author contributions

Conceptualization, I.S.P.; methodology, M.A.H.; investigation, M.A.H. & M.S.H.; writing-original draft preparation, M.A.H.; writing-review and editing, M.A.H.; supervision, I.S.P.

## Acknowledgement

Thanks to Rufaida BioMeds for providing their lab facilities

## Conflicts of interest

None to declare

## Institutional review board statement

Not applicable

## Data availability statement

All relevant data are within the manuscript and its Supporting Information files.

## Notes

### Competing Interest Statement

The authors have declared no competing interest.

## References

1. Barnes, D.E. and K. Yaffe, The projected effect of risk factor reduction on Alzheimer’s disease prevalence. Lancet Neurol, 2011. 10(9): p. 819–28.

2. Sun, X., W.D. Chen, and Y.D. Wang, *beta-Amyloid: the key peptide in the pathogenesis of Alzheimer’s disease*. Front Pharmacol, 2015. 6: p. 221.

3. Chen, G.F., et al., Amyloid beta: structure, biology and structure-based therapeutic development. Acta Pharmacol Sin, 2017. 38(9): p. 1205–1235.

4. Terry, R.D., N.K. Gonatas, and M. Weiss, Ultrastructural Studies in Alzheimer’s Presenile Dementia. Am J Pathol, 1964. 44: p. 269–97.

5. Duyckaerts, C., B. Delatour, and M.C. Potier, Classification and basic pathology of Alzheimer disease. Acta Neuropathol, 2009. 118(1): p. 5–36.

6. McLean, C.A., et al., Soluble pool of Abeta amyloid as a determinant of severity of neurodegeneration in Alzheimer’s disease. Ann Neurol, 1999. 46(6): p. 860–6.

7. Cleary, J.P., et al., Natural oligomers of the amyloid-beta protein specifically disrupt cognitive function. Nat Neurosci, 2005. 8(1): p. 79–84.

8. Haass, C. and D.J. Selkoe, Soluble protein oligomers in neurodegeneration: lessons from the Alzheimer’s amyloid beta-peptide. Nat Rev Mol Cell Biol, 2007. 8(2): p. 101–12.

9. De, S., et al., Different soluble aggregates of Abeta42 can give rise to cellular toxicity through different mechanisms. Nat Commun, 2019. 10(1): p. 1541.

10. Shahnawaz, M., T. Bilkis, and I.S. Park, Amyloid beta cytotoxicity is enhanced or reduced depending on formation of amyloid beta oligomeric forms. Biotechnol Lett, 2021. 43(1): p. 165–175.

11. Wells, C., et al., The role of amyloid oligomers in neurodegenerative pathologies. Int J Biol Macromol, 2021. 181: p. 582–604.

12. Kienlen-Campard, P., et al., Intracellular amyloid-beta 1-42, but not extracellular soluble amyloid-beta peptides, induces neuronal apoptosis. J Biol Chem, 2002. 277(18): p. 15666–70.

13. Wong, P.T., et al., Amyloid-beta membrane binding and permeabilization are distinct processes influenced separately by membrane charge and fluidity. J Mol Biol, 2009. 386(1): p. 81–96.

14. Chafekar, S.M., F. Baas, and W. Scheper, Oligomer-specific Abeta toxicity in cell models is mediated by selective uptake. Biochim Biophys Acta, 2008. 1782(9): p. 523–31.

15. Haque, M.A., et al., Evidence for a Strong Relationship between the Cytotoxicity and Intracellular Location of beta-Amyloid. Life (Basel), 2022. 12(4).

16. Spencer, J.P., Flavonoids: modulators of brain function? Br J Nutr, 2008. 99 **E Suppl 1**: p. ES60–77.

17. Bagchi, D., et al., Acute and chronic stress-induced oxidative gastrointestinal mucosal injury in rats and protection by bismuth subsalicylate. Mol Cell Biochem, 1999. 196(1-2): p. 109–16.

18. Yamamoto, Y. and R.B. Gaynor, Therapeutic potential of inhibition of the NF-kappaB pathway in the treatment of inflammation and cancer. J Clin Invest, 2001. 107(2): p. 135–42.

19. Cushnie, T.P. and A.J. Lamb, Antimicrobial activity of flavonoids. Int J Antimicrob Agents, 2005. 26(5): p. 343–56.

20. Commenges, D., et al., Intake of flavonoids and risk of dementia. Eur J Epidemiol, 2000. 16(4): p. 357–63.

21. Beking, K. and A. Vieira, Flavonoid intake and disability-adjusted life years due to Alzheimer’s and related dementias: a population-based study involving twenty-three developed countries. Public Health Nutr, 2010. 13(9): p. 1403–9.

22. Yao, Z., K. Drieu, and V. Papadopoulos, The Ginkgo biloba extract EGb 761 rescues the PC12 neuronal cells from beta-amyloid-induced cell death by inhibiting the formation of beta-amyloid-derived diffusible neurotoxic ligands. Brain Res, 2001. 889(1-2): p. 181–90.

23. Luo, Y., et al., Inhibition of amyloid-beta aggregation and caspase-3 activation by the Ginkgo biloba extract EGb761. Proc Natl Acad Sci U S A, 2002. 99(19): p. 12197–202.

24. Ono, K., et al., Effects of grape seed-derived polyphenols on amyloid beta-protein self-assembly and cytotoxicity. J Biol Chem, 2008. 283(47): p. 32176–87.

25. Kim, J.K., et al., Protective effects of kaempferol (3,4’,5,7-tetrahydroxyflavone) against amyloid beta peptide (Abeta)-induced neurotoxicity in ICR mice. Biosci Biotechnol Biochem, 2010. 74(2): p. 397–401.

26. Rigacci, S., et al., Abeta(1-42) aggregates into non-toxic amyloid assemblies in the presence of the natural polyphenol oleuropein aglycon. Curr Alzheimer Res, 2011. 8(8): p. 841–52.

27. Lu, J.H., et al., Baicalein inhibits formation of alpha-synuclein oligomers within living cells and prevents Abeta peptide fibrillation and oligomerisation. Chembiochem, 2011. 12(4): p. 615–24.

28. Qin, X.Y., Y. Cheng, and L.C. Yu, Potential protection of green tea polyphenols against intracellular amyloid beta-induced toxicity on primary cultured prefrontal cortical neurons of rats. Neurosci Lett, 2012. 513(2): p. 170–3.

29. Ramassamy, C., Emerging role of polyphenolic compounds in the treatment of neurodegenerative diseases: a review of their intracellular targets. Eur J Pharmacol, 2006. 545(1): p. 51–64.

30. Feng, Y., et al., Resveratrol inhibits beta-amyloid oligomeric cytotoxicity but does not prevent oligomer formation. Neurotoxicology, 2009. 30(6): p. 986–95.

31. Zhu, J.T., et al., Flavonoids possess neuroprotective effects on cultured pheochromocytoma PC12 cells: a comparison of different flavonoids in activating estrogenic effect and in preventing beta-amyloid-induced cell death. J Agric Food Chem, 2007. 55(6): p. 2438–45.

